# Uncursing winner’s curse: on-line monitoring of directed evolution convergence

**DOI:** 10.1101/2023.01.03.522172

**Authors:** Takahiro Nemoto, Tommaso Ocari, Arthur Planul, Muge Tekinsoy, Emilia A. Zin, Deniz Dalkara, Ulisse Ferrari

## Abstract

Directed evolution (DE) is a versatile protein-engineering strategy, successfully applied to a range of proteins, including enzymes, antibodies, and viral vectors. However, DE can be time-consuming and costly, as it typically requires many rounds of selection to identify desired mutants. Next-generation sequencing allows monitoring of millions of variants during DE and can be leveraged to reduce the number of selection rounds. Unfortunately the noisy nature of the sequencing data impedes the estimation of the performance of individual variants. Here, we propose ACIDES that combines statistical inference and in-silico simulations to improve performance estimation in DE by providing accurate statistical scores. We tested ACIDES first on a novel random-peptide-insertion experiment and then on several public datasets from DE of viral vectors and phage-display. ACIDES allows experimentalists to reliably estimate variant performance *on the fly* and can aid protein engineering pipelines in a range of applications, including gene therapy.

## INTRODUCTION

Directed evolution (DE) [1–3] is a versatile protein engineering strategy to conceive and optimize proteins like enzymes [4–6], antibodies [7, 8] or viral vectors for gene therapy [9–15], culminating in the Nobel Prize in Chemistry 2018 [16]. DE starts from a massive library of random mutants, screens it against a given task over multiple rounds and searches for the variants with the highest performance. As the iteration continues, the best performing variants get enriched and emerge from the bulk, while ineffective ones are instead weeded out. Nowadays, we can rely on next generation sequencing (NGS) [17, 18] to sample millions of variants within the library and monitor their concentrations over multiple rounds or time-points. In this approach, the enrichment of the screened variants is measured to rank the variants depending on their performance. In a similar flavor, Deep mutational scanning (DMS) experiments [19–21] combine extensive mutagenesis with NGS to study the properties of proteins [22–26], promotors [27, 28], small nucleolar RNA [29], or other amino-acid chains. It uses similar techniques to DE and requires similar analysis. The approach presented here can be applied to both DE and DMS experiments, and focus on their common issues and needs.

The analysis of NGS data of multiple selection rounds presents several difficulties. First, variants need to be robustly scored based on their enrichment rates, so-called selectivities [30, 31]. This task is complicated by the large noise in the NGS counts introduced by, for example, polymerase chain reaction (PCR) amplification or bacterial cloning, during amplicon preparations [32–34]. This noise needs to be taken into account in the analysis. Second, in order to rank the variants and to identify the best performing ones, the score should come with a precise estimation of its statistical error. As a consequence of the noise in the counts, some irrelevant variants might appear to be highly enriched (winner’s curse). This would be anticipated if properly estimated credibility scores are available. Third, when running DE over multiple rounds, it is hard to know when to end the experiment: performing too few rounds could lead to selection of weak variants, not representative of their true ranking. On the other hand, performing too many rounds is costly, time-consuming and even ethically questionable when working with *in-vivo* selections [14, 35]. Similarly, it would be useful to understand the best NGS depth for a given experiment, as deepening the NGS by increasing reads results in better data, but adds an extra expense to the experiment.

In order to account for these issues and needs, we present ACIDES, Accurate Confidence Intervals for Directed Evolution Scores, a computational method to empower the analysis of DE and DMS experiments. We focus on screening experiments on highly diverse libraries where massive NGS data are collected over multiple rounds or multiple time-points (Fig. 1A). Our goal is to develop a method to extract maximal information from noisy NGS data, and allows for scoring and ranking variants with accurate statistical confidence scores. Our approach can be applied to different kinds of experiments, such as *in-vivo* DE [13, 14, 36], and DMS of phage-display [23, 30, 37], yeast two-hybrid [23] and small nucleolar RNA [29] experiments. It is possible to apply ACIDES either *a posteriori* over data collected previously, or along the course of the experiment as soon as the NGS data become available. The latter strategy allows for monitoring the selection convergence *on the fly*, and to understand when the experiment can be ended. In this way, ACIDES can be integrated into protein engineering pipelines as well as studies of protein function using mutagenesis. The tutorial for using ACIDES, along with an executable code in Python, will be available in GitHub upon publication of this manuscript.

**FIG. 1.**
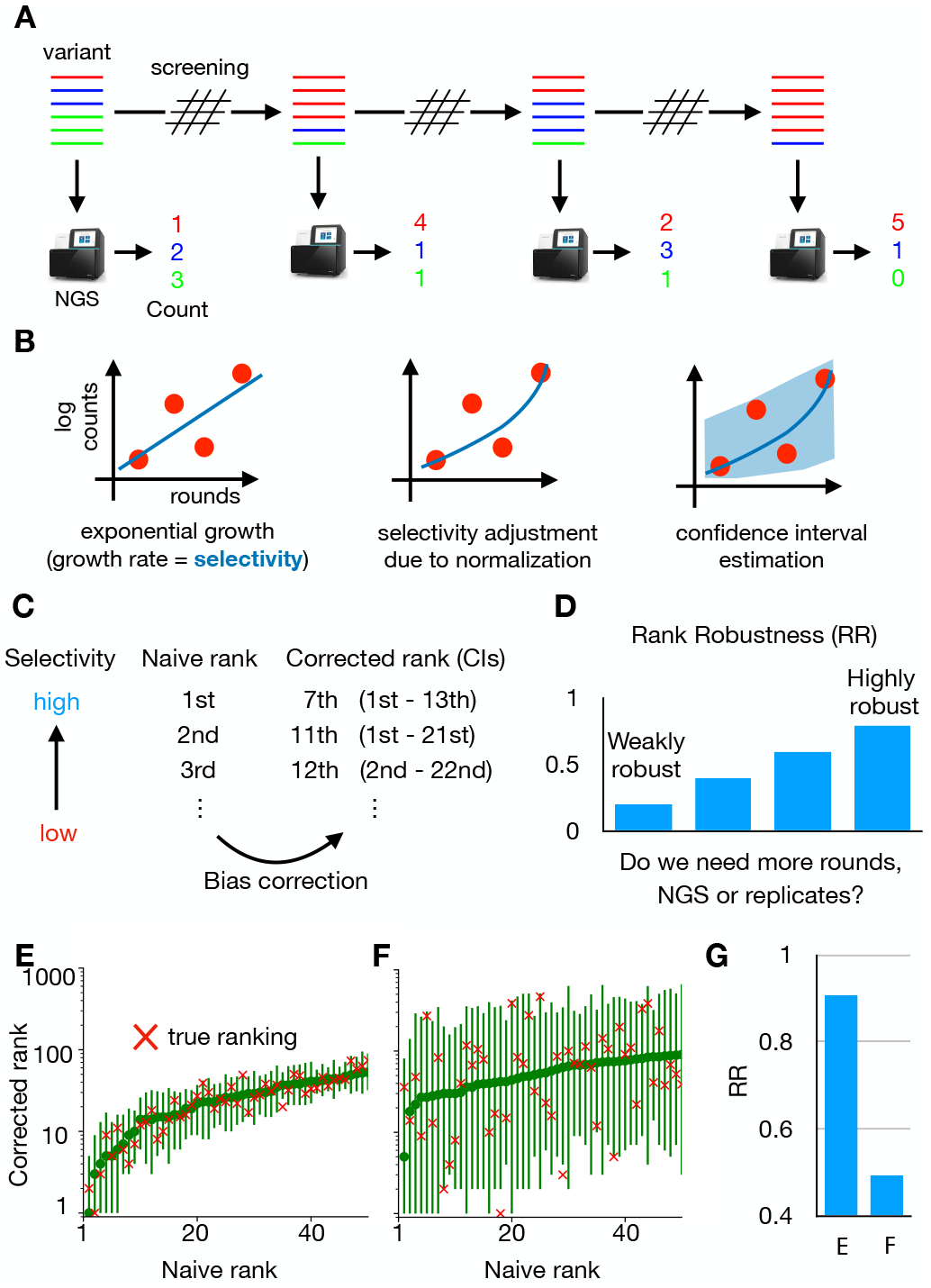
ACIDES framework. (A) We consider directed evolution (DE) experiments, where protein variants are screened over multiple rounds, and massive NGS datasets are collected. (B) From the obtained count data, we estimate a score (selectivity) for each variant. The higher the score, the better the variant for the task. Each score is estimated together with its 95%-confidence interval (CI). (C) Sorting the scores of all variants in descending order, we obtain a variant rank (naive rank). Due to statistical errors in the scores, the obtained rank is biased in general. To correct for this, using *in-silico* simulations based on the CIs of the scores, we reestimate the rank with 95%-CI (corrected rank). (D) From the obtained corrected rank, we compute Rank Robustness (RR). RR represents the percentage of the top 50 variants identified in the naive rank that also appear in the top 50 of the corrected rank. (E,F) Examples of rank graphs for two synthetic datasets with different depths of NGS (per round) and numbers of unique variants (respectively, E: 10^7^, 5 ×10^4^; F: 10^6^ and 10^6^). The true rank is shown as red crosses. In both cases, most red crosses are within the 95%-CI of the corrected rank. (G) RR for the two synthetic datasets. Note that RR multiplied by 50 (E: ∼45.3; F: ∼24.6) roughly provides the number of the correct top-50 sequences, which are 46 and 23, respectively. (See Fig.s S3 and S4 for more systematic comparison).

## RESULTS

The first step of ACIDES estimates the selectivity of each individual variant present in the dataset (Fig. 1B) and its 95% confidence interval (95%-CI). In this study the term selectivity means the rate at which each variant increases its concentration with respect to the others. More precisely, we assume an exponential growth as 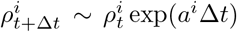, where 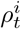 is the concentration of variant *i* at time *t*, and *a*^*i*^ is its selectivity. Compared with previous methods [19–24, 27–31, 38], our approach combines a robust inference framework (maximum likelihood estimation) with a better quantification of the NGS sampling noise [32–34]. For this scope, our approach benefits from a negative binomial distribution [39–42] (Fig. S1) in which the variance of the noise is overdispersed and grows as *λ* + *λ*^2−*α*^*/β*. Here *λ* is the expected mean count, and *α, β* are parameters to be inferred (Materials and Methods). Using novel data from a plasmid library, we observed that our negative binomial model realizes a 50- to 70-fold improvement over the Poisson model in the predictive ability of the NGS sampling noise (Fig. S1). The second step of ACIDES uses the estimation of the selectivities and their statistical errors to rank the variants. The rank obtained by sorting the selectivities in descending order (naive rank) is biased due to statistical fluctuations of the selectivities. We correct this bias using *in-silico* simulations (Fig. 1C). The third and last step of ACIDES uses simulations to quantify a Rank Robustness (RR), a measure of the quality of the selection convergence (Fig. 1D). Specifically, RR is the ratio at which the top-50 variants in the naive rank are correctly identified (Materials and Methods). RR ranges from 0 to 1: a low value points out that the variants have not been selected enough, and therefore calls for the necessity to perform more rounds, deeper NGS sampling or possibly more replicates. Conversely, a large value confirms that the selection has properly converged, and suggests that the experiment can be ended without performing additional experimental steps.

Before focusing on experimental data, we apply ACIDES to two synthetic datasets (Materials and Methods) describing two opposite scenarios (See Fig.s S3 and S4for more systematic comparison): data-rich case (more NGS reads with fewer unique variants) and data-poor case (less NGS reads with more considered variants). In the data-rich case, we first verify that our method reaches high performance in recovering the ground-truth values of the selectivities (*R*^2^ ≃ 0.92, Fig.S3) in a teacher-student settings. In this first case, selection convergence is reached and the different variants can be robustly ranked (Fig. 1E). In the data-poor case, instead, CI-bars are king is uncertain (Fig. 1F). Consistently, the estimates RRs are high and low for, respectively, the data-rich and poor examples (Fig. 1G). Note that, once multiplied by 50 RR roughly provides the number of the correct top-50 variants in both cases (caption of Fig. 1G). Furthermore, we observe that most true rank values (red crosses fall within the 95%-CI in both examples. These that our approach can quantify statistical errors even in the data-poor regime (See Fig. S4 for more systematic comparison).

### Analysis of directed evolution and deep mutational scanning experiments

In order to showcase ACIDES, we apply it to several screening datasets, where various proteins (and one RNA molecule) are screened using different experimental techniques (Table I). Specifically, we consider three phage-display screening experiments targeting different proteins, such as the breast cancer type 1 susceptibility protein (BRCA1) for Data-A, human yes-associated protein 65 (hYAP65) for Data-F and immunoglobulin heavy chain (IgH) for Data-G, two *in-vivo* DEs of adenoassociated virus type 2 (AAV2) vectors targeting canine eyes for Data-C and murine lungs for Data-D, a multiplexed yeast two-hybrid assay targeting BRCA1 for Data-B and a yeast competitive growth screen measuring the fitness of mutant U3 gene for Data-E. For each of these experiments, we rank variants (naive rank) and compute the confidence interval of their ranks (corrected rank in Fig. 2A-G). The degree of convergence of the selection is quantified by RR (2H). When technical replicates are available (Data-A and Data-B), we compute RR over all of them and obtained consistent results (shown by the small error-bars in Fig. 2H).

**TABLE I.** Next generation sequencing datasets of directed evolution experiments

| Label | Experiment | Target | Time-points | Reads/round | # of variants | Replicates | Ref |
| --- | --- | --- | --- | --- | --- | --- | --- |
| A | Phage display | BRCA1 | T0 → T5 | 13.6 M | 35 k | 2 × 3 | Starita 2015 [23] |
| B | Yeast two-hybrid | BRCA1 | T0 → T3 | 13.5 M | 27 k | 2 × 3 | Starita 2015 [23] |
| C | *in-vivo* DE (dog eye) | AAV2-7mer | T1 → T5 | 17 M | 5 k | 1 × 1 | Byrne 2018 [36] |
| D | *in-vivo* DE (murine lung) | AAV2-7mer | T0 → T5 | 6.2 M | 0.5 k | 1 × 1 | Korbelin 2016 [13] |
| E | Yeast competitive growth | U3 snoRNA | T0 → T4 | 8 M | 24 k | 2 × 1 | Puchta 2016 [29] |
| F | Phage display | hYAP65 WW | T0 → T3 | 5 M | 470 k | 2 × 1 | Araya 2012 [30] |
| G | Phage display | Ab IgH | T1 → T3 | 0.1 M | 29 k | 1 × 1 | Boyer 2016 [37] |

**FIG. 2.**
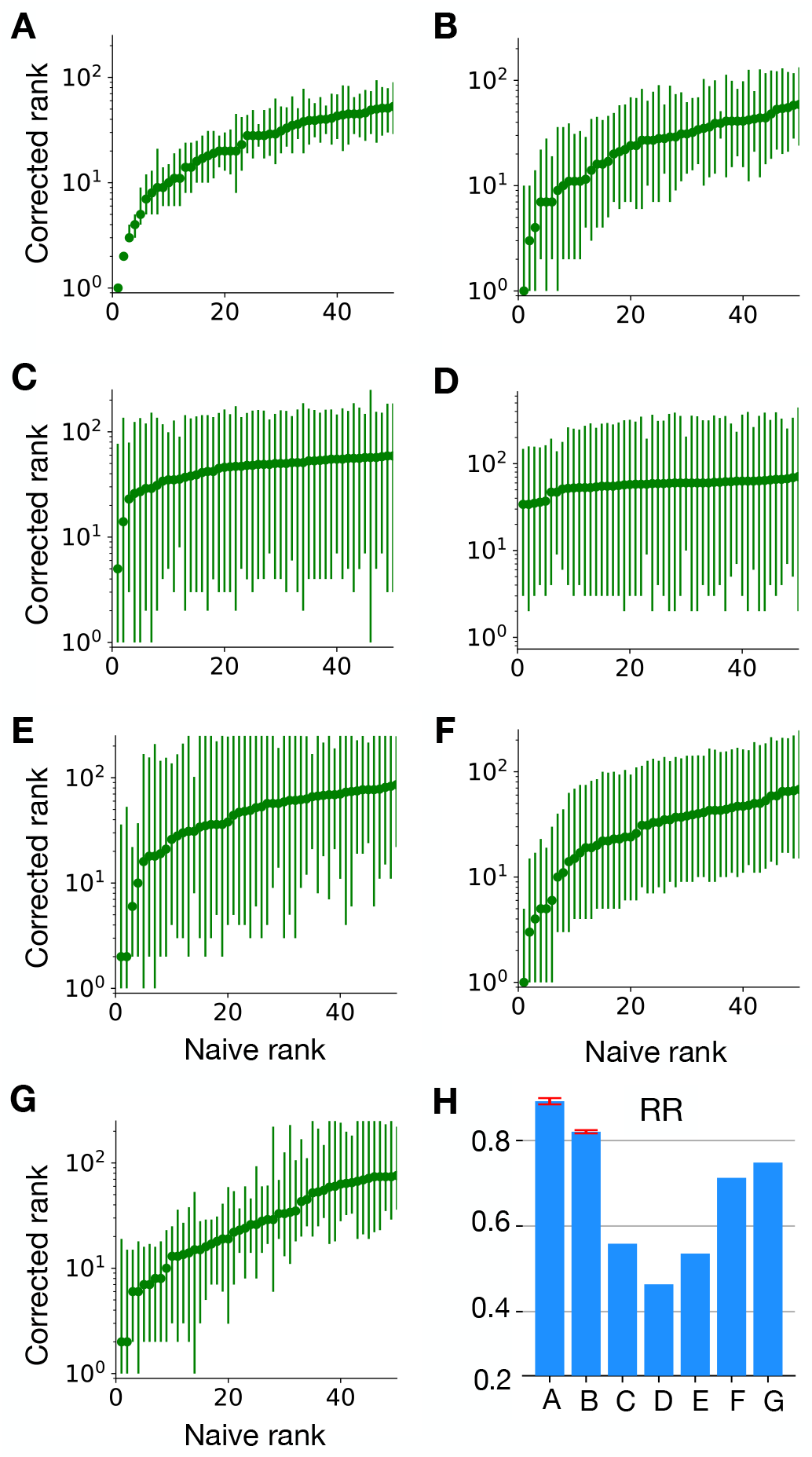
Rank graph for various experimental datasets. The panel labels A-G correspond to the experiments listed in table I. (H) Rank robustness (RR) for each experiment. When technical replicates are available (Data-A and -B), the mean and standard deviation are shown.

We classify the observed RRs into three groups depending on the quality of the selection convergence: high (Data-A and Data-B), intermediate (Data-F and Data-G), and low (Data-C, Data-D and Data-E) convergence groups. The high group seems to behave similarly to the data-rich synthetic data in Fig. 1E. Consistently, RR, NGS depth and the number of unique variants are indeed of the same order (Table I). In these cases, the obtained naive rank is robust, as indicated by the value of RR (RR *>* 0.8). In the intermediate group, the value of RR ranges between 0.6 and 0.8. The experimental techniques used in these datasets are similar to those in the high group, but the NGS depths (or the numbers of unique variants) are smaller (or larger), which could be the reason why they result in lower RRs. The low group suffers from the noise in the data. In Data-C and Data-D, the numbers of unique variants are lower than those of Data-A and Data-B, which would normally help these datasets with having higher RR, given the same NGS depth. As this is not the case, we see that some experiments are *intrinsically* more difficult than the others, *i*.*e*., in-vivo DE (Data-C and Data-D) and RNA based screening (Data-E) will result in lower RRs than the other experiments if the NGS depth and number of variants are similar.

In datasets with low RRs, some variants seem to perform better than the others, but the difference between their scores is marginal compared with their statistical errors. This means that we cannot distinguish if the obtained variants are selected because of their ability to perform the task (fitness) or just there due to noise. In these cases, experimentalists have two possibilities: (i) based on the noisy identified variants, perform further tests in addition to DE [13, 14], as for example, study infective ability of viral vectors using single-cell RNA-seq [43]. Or (ii) increase the quality of the datasets, by performing further selection rounds, increasing NGS depths, or replicating the experiments under the same conditions. This second possibility is explored in the next section. Overall our rank-analysis of the different experiments shows how our approach can provide an overview of the selection convergence, informing about the state of the experiment and eventually pointing out the necessity of more experimental efforts.

### Integration into the experimental pipeline

Noise in experimental data can be reduced by performing additional selection rounds involving experiments, but in general these are expensive, time-consuming and, in case of experiments involving animal use, ethically problematic [35]. For these reasons, it is important to choose accurately the number of rounds and the NGS depth. For this scope, ACIDES can be integrated into experimental pipelines to obtain an overview on how RR depends on these factors. This is to help experimentalists with making informed decisions about additional experimental efforts.

ACIDES can estimate RR after each selection round (or any time new data become available). This allows us to examine the data’s behavior and to quantify the degree of convergence in terms of the selection rounds. Similarly, for each round, ACIDES can be run on down-sampled NGS data to compute RR with smaller NGS depth (Materials and Methods). Using these two techniques, we monitor the need for more selection rounds or deeper NGS: a slow increase of RR (or no change in RR) upon improving data-quality implies that convergence is reached and suggests that the experiment can be ended. If, on the other hand, RR increases rapidly when improving the rounds and/or NGS depth, it is probably worth making further experimental efforts.

In order to showcase our approach, we study how RR depends on the number of screening rounds and NGS depth in previous experiments. We start by measuring RR in Data-A for different NGS depths. 95%-CI on corrected ranks gets larger as the NGS depth becomes smaller (Fig.3A). At 1% NGS depth, the variant ordering seems largely unreliable: RR is smaller than 0.5 (Fig.3B). Importantly, RR does not decrease smoothly as the NGS depth decreases, but it remains roughly constant at the beginning, and falls only at a very small NGS depth. This result suggests that the actual NGS depth of this experiment largely exceeds what was necessary (10% of the depth would have been sufficient). Next, we quantify how RR depends on both the number of performed rounds and NGS depth (Fig.3C). RR grows from 0.28 (3 performed rounds with 1% NGS depth) to 0.88 (6 performed rounds with 100% NGS depth). Saturation of RR seems to be observed for RR *>* 0.7, which corresponds to 5 performed rounds with the NGS depth larger than 20%, or 4 performed rounds with the NGS depth larger than 40%. This again indicates that the experiment could have been stopped earlier (less rounds and/or lower sequence coverage) without much affecting the outcome. Note that different datasets show different behaviors. For Data-E more selection rounds with a higher number of NGS reads is expected to improve RR, while for Data-B they seem to have just reached the saturation point (Fig.3D).

Overall these results show how our approach can be implemented along experimental pipelines. By estimating RR while collecting new data, we can understand if we should continue/stop adding more rounds or increasing NGS depth. This could avoid unnecessary, costly and time-consuming experimental efforts. Similar analyses can be done on the number of replicate experiments (Fig.S6).

### Comparison with previous work

We compare the performance of ACIDES with *Enrich2*, the state of the art for estimating variant scores (selectivities) [31]. *Enrich2* is based on a weighted linear fitting of the log-count change along rounds, and the first step of ACIDES should be seen as an upgrade for this fitting. In order to compare these two approaches we take advantage of replicate datasets. We first investigate if the scores in each method are consistent over replicates. For this, we plot the scores obtained from one replicate against the scores obtained from the other (Fig. 4A, B). The correlation between replicates is estimated using the coefficient of determination (R^2^). The correlation quantifies the quality of the method, as higher (or lower) correlations imply that the estimated scores are more (or less) robust and fewer (or more) replicates are needed to obtain reliable results. The figure shows that ACIDES outperforms *Enrich2*. Next, we test how the comparison depends on the data quality. To this goal, we systematically select a set of variants based on the magnitude of predicted statistical score-errors (Materials and Methods). (Smaller/larger sets include variants with smaller/larger predicted statistical errors.) For each set, we measure the correlation between two replicates as in (Fig. 4A, B), and plot it as a function of the set size (Fig. 4C). We observe ACIDES’s correlation becomes more dominant as the set size decreases, suggesting the better quality of both the estimated scores and statistical errors. In order to generalize these results, we perform the comparison for all possible 12 pairs of technical replicates in Data-A and Data-B (Table I). In all cases our approach outperforms the competitor (Fig. 4D). We also perform an additional test to quantify the consistency of the predicted statistical errors (Supp. Fig. S7).

**FIG. 4.**
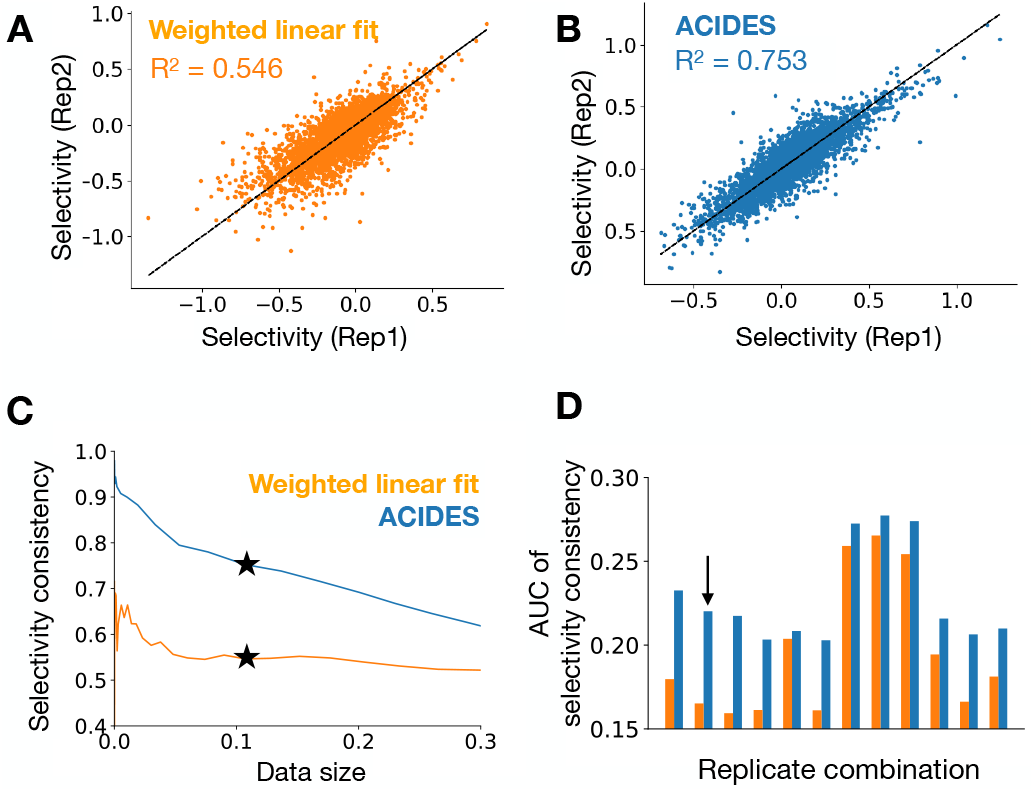
Comparison of our approach with the state of the art. Using technical replicates in Data-A, we compare ACIDES with a weighted linear least squares method (*Enrich2*) [31]. For both methods (*Enrich2* (A) and ACIDES (B)), the inferred selectivities from one replicate are plotted against the selectivities in the other replicate. The coefficient of determination (*R*^2^), which quantifies the consistency between two replicates, is also shown. (C) We next examine how the comparison in the panels A and B depend on data quality. We consider a set of variants in which the estimated statistical errors (Materials and Methods) are smaller than a given threshold. Varying this threshold, sets of variants are systematical selected, where larger/smaller sets include variants with larger/smaller estimated statistical errors. For each set, we estimate *R*^2^ between two replicates, and plot it as a function of the set size. The panel A and B correspond to the stars *\** in C (data size 0.11). (D) In order to test both methods more systematically, we perform the same analysis (as those in the panels A-C) for all possible 12 combinations of technical replicates in Data-A and Data-B. We define the area under curve of *R*^2^ (in the panel C) and plot it for these combinations (D). Our method systematically outperforms the weighted linear fitting method. The replicate combination used for the panels A-C is indicated by the arrow in the panel D.

## DISCUSSION

In this work we have presented ACIDES, a method to quantify DE and DMS selectivities (fitness), rank variants with accurate credibility scores and measure the degree of experimental convergence. ACIDES can be used *on the fly* to offer an overview of the progress of selection experiments, which would help experimentalists with making informed decisions on whether new experimental efforts are needed. In this way, ACIDES can save significant experimental time and resources. We have applied ACIDES to several DE and DMS datasets where a number of different target proteins and one set of target RNA molecules have been screened using different experimental protocols. The heterogeneity of these datasets shows that ACIDES is a method of general use, applicable to many different experiments.

The first step of ACIDES estimates the score (selectivity) of each observed variant. This is a necessary step, and several alternative methods have been proposed in the past. In many applications, such scores are computed as the variant enrichment that is defined as the logarithmic ratio between the variant frequencies in the last and second to last round [13] or between the last and first round [14, 19, 20, 22, 27, 38]. These approaches thus make use of data from only two rounds and disregard all the others. For this reason, this strategy is suboptimal and may lead to noisy score estimations. A more sophisticated approach that uses all the data consists in inferring the slope of a linear line fitted to the log-frequencies of variants over all the screening rounds/time points [23, 24, 28, 30]. This method gives the same importance to log-frequencies in all the rounds. Yet as variant counts in the first rounds are typically small and noisy, assuming the same weight on them could result in an overfitting. To fix this effect, Enrich2 [31] uses the variance of the count data - estimated via a Poisson distribution assumption - as the weights in a linear least squares fitting. ACIDES’ first step comes with a three-fold improvement over this last approach. First, instead of relying on the linear least squares fitting, we estimate the score by log-likelihood maximization. A major improvement happens for variants whose log-frequencies do not grow linearly with the rounds, and a simple linear weighted fit may struggle in identifying the correct slope. This is particularly visible in the bulk variants with intermediate scores (Fig.4 A, B). Secondly, instead of a simple exponential growth of the counts, we included a softmax non-linear function (Materials and Methods), where the denominator is inferred from data [44]. This change improves the score estimation when the wildtype (if any) and/or few variants have a large fraction of the total counts and bend the exponential growth of the log-frequencies. Lastly, ACIDES uses a negative binomial distribution to model the count variability [39–42] This distribution accounts for the large dispersion of next generation sequencing data [32–34] far better than the Poisson distribution (Fig. S1). Additionally, the negative binomial loss in the likelihood maximization allows us to better estimate statistical errors for the inferred scores. Thanks to all these improvements, our approach realizes a more robust and accurate estimation of the variant scores and outperforms the previous method (Fig. 4).

In case of noisy data, the estimated scores of variants come with statistical errors. This means that the rank obtained from the scores (naive rank in our figures) is in general biased: top ranked variants are overvalued, and vice-versa. This simple statistical effect was not taken into account in previous analyses related to DE and DMS experiments. The second step of ACIDES uses a bootstrap method to account for the bias and recover both the corrected rank and its 95%-CI. The deviation between this 95%-CI and the naive rank shows us how much we can trust the naive rank. To quantify it, as a third step of ACIDES, we introduce RR that describes how many of the top-50 variants in the naive rank are correct. RR measures how stable and robust are the ranks of the variant selectivities. As such, it quantifies the degree of convergence of the experimental selection, providing an insightful overview of the state of the experiment.

Although ACIDES demonstrates advantages over the other methods, it has several limitations that may be addressed in the future. First of all, ACIDES does not account for changes in the selection pressure over rounds. This can potentially be included, but has not been done here, as the selection pressure is constant in most datasets we analyzed in this article. Second, ACIDES uses a negative binomial model to describe the dispersion of count data by assuming that the count variance depends only on the frequency of the variant. Although this assumption proves useful to describe NGS count errors (Fig. S1) and is used elsewhere [42], it is possible that dispersions induced by a sequence-dependent procedure, such as error-prone PCR [14, 36, 45], may not be taken into account by our method (Note that Data-C includes an error-prone PCR after the third round of selections, indicating that the estimated results for Data-C may contain biases). We would need to analyze more data from DE experiments using error-prone libraries to address this question. Third, statistical errors due to the replicates that do not share the same initial library cannot be described by ACIDES, provided that the model is only trained on a single series of screening rounds. To account for this, we would need a framework that generalizes ACIDES for different sources of variability.

Finally, using machine learning techniques, several studies have aimed at estimating selectivities from the amino-acid sequences of variants. Most of these methods rely on supervised algorithms, which are trained to predict the selectivity (output) from the sequence of a variant (input) [45–53]. Because the performance of these methods depends on how the selectivity is estimated from data, ACIDES can potentially be incorporated in their pipelines to improve the overall performance. We leave such analysis for future developments. Other methods use instead unsupervised approaches to predict selectivities from the sequences of variants [44, 54–57]. Even if these methods do not use any sequence scores for their training, they often use it to validate and/or test the model. Our approach would therefore be useful also in these cases.

## METHOD

### Library preparation for Fig.S1

To demonstrate that our negative binomial likelihood approach outperforms the Poisson counterpart, we conducted the following experiment: We inserted random 21 nucleotide oligomers into a RepCap plasmid containing adenoassociated virus 2 (AAV2) cap gene using previously described methods [58]. The plasmid library obtained was deep sequenced following generation of amplicons corresponding to the 7mer insertion region. Since the 21 nucleotides are randomly and independently generated, we can use a position weight matrix model to predict the frequency of each variant in the sample. Based on this property, the performance of the two models are examined as shown in Fig.S1.

### Model

We propose ACIDES for analyzing selection data in DE and DMS. Here the mathematical model is described in detail. For a given series of samples over screening rounds, we perform NGS and denote by 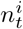 the obtained count of the *i*-th variant (*i* = 1, 2, …, *M*) at round (time-point) *t* ∈*T*. We denote by *N*_*t*_ the total count 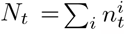 at *t*. For each sample, we define 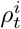 as the *expected* value of frequency of the *i*-th variant at *t*. (Note that “expected” means that 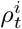 itself does not fluctuate due to the noise in the experiment.) For each variant, an initial frequency 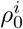 and a growth rate *a*^*i*^ are assigned, by which the expected frequency is computed as

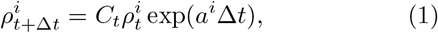

where Δ*t* is the round- (or time-) difference between two consecutive NGSs. *C*_*t*_ is a normalization constant, defined as 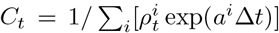. We call this model ((1)) an exponential model.

We use a negative binomial distribution 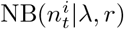 with two parameters *λ* and *r* to model the noise distribution of counts 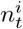. Here *λ* is the expected value of count 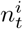 given as 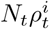, while *r* is the dispersion parameter that describes the deviation of the negative binomial distribution from the Poisson distribution. (The negative binomial distribution is a generalization of the Poisson distribution with a variance equal to *λ*(1 + *λ/r*): the Poisson distribution is recovered in the large *r* limit.) Here, based on Fig. S1 and [42], we assume *r* is a power-law function of *λ*: *r*(*λ*) = *βλ*^*α*^ (with *α, β >* 0), where *α* and *β* are parameters that are common for all the variants in the experiment. (The variance is thus *λ* + *λ*^2−*α*^*/β*.) Model parameters *α, β* as well as 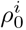, *a*^*i*^ (*i* = 1, 2, …, *M*) are inferred from the count data 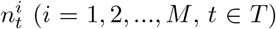 by maximizing the following likelihood function:

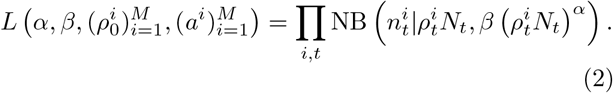

The 95%-CIs of the estimated parameters are computed from the curvature of the log-likelihood function at the maximum.

### Synthetic data

Synthetic count data 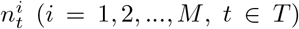 are generated from the model ((2)) for a given parameter set *α, β*, 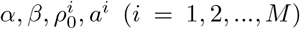, *a*^*i*^ (*i* = 1, 2, …, *M*). For Fig.1, we use *α, β* = 0.69, 0.8 with 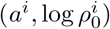 generated from the normal distribution with the expected values (−1, 1) and the standard deviations (0.25, 1). (*M, N*_*t*_) are (5 ×10^4^, 10^7^) for the data-rich case (Fig.1E) and (10^6^, 10^6^) for the data-poor one (Fig.1F).

### Model inference

To maximize the likelihood function, we develop a two-step algorithm. The first step infers (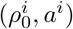, *a*^*i*^), while the second (*α, β*) and then we iterate the two steps until convergence is reached. All inferences are done with a gradient descent algorithm, and to reach convergence 10-30 iterations are usually sufficient. The first step is itself iterative, and loops between the inference of (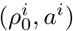, *a*^*i*^) and *C*_*t*_ by treating *C*_*t*_ as a parameter. Here we also introduce a gauge choice because of the redundancy between 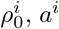, *a*^*i*^ and *C*_*t*_ (the caption of Fig. S2 for more details). In the second step, the inference of (*α, β*) with a straightforward gradient method produces a bias (Fig. S2E). In order to correct this, at each iteration the algorithm adopts a teacher-student framework, runs a simulation of the count data with the current parameters to obtain an estimation of the bias, which is then used to correct the real inference and update the parameters.

In order to reduce computational time and to increase the stability of the algorithm, we first run the inference algorithm on a subset of variants to estimate *α, β*. We then compute (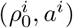, *a*^*i*^) of the excluded variants using the estimated *α, β*. For this subset, we use the variants that satisfy the following two criterions: (i) their counts are larger than 0 more than twice in the selection rounds and (ii) whose total NGS count (as summed over all the rounds) is above a threshold. We set this threshold to 100 for all the datasets except for Data-C -D, where 10000 is used. This is because the noise in these experiments is larger than the others. Results are stable by changing the threshold value (Fig. S2F).

### Simulated rank and rank robustness (RR)

Using the standard deviations *δa*^*i*^ (*i* = 1, …, *M*) of estimated scores *a*^*i*^, we discard the variants with higher estimated errors. We keep 5000 variants for further analysis and denote by *A* their indices. We then rearrange the variant index in *A* in descending order of *a*^*i*^ to define a *naive rank* (the *x*-axis of Fig 2A-G). To obtain a *corrected rank* (the *y*-axis of Fig 2A-G), we first generate synthetic scores using the normal distribution with the expected value (*a*^*i*^)_*i*∈*A*_ and the standard deviation (*δa*^*i*^)_*i*∈*A*_. Based on the generated scores, we rearrange the variant index in descending order and define a synthetic naive rank. Repeating this estimation 3000 times, we then compute the median and 95%-CI of the obtained synthetic naive ranks. This 95%-CI is defined as the corrected rank.

To estimate RR, we compare the top-50 variants in the naive rank and each synthetic naive rank. We count the number of overlaps between them and average it over the 3000 estimations. RR is computed by dividing the obtained overlap by 50.

### NGS-Downsampling for Fig.3

To obtain downsampled count data 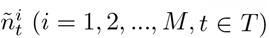 by a factor *E*, we sample synthetic data from the likelihood function ((2)) with a reduced number of the total counts *EN*_*t*_ (*t*∈ *T*) and with the estimated parameters 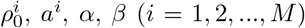, *a*^*i*^, *α, β* (*i* = 1, 2, …, *M*). To obtain a downsampled RR in Fig. 3, we first re-estimate *a*^*i*^ (*i* = 1, 2, …, *M*) from 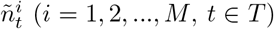 using the values of (*α, β*) that are already known, and then perform the estimation of RR described above. Using the synthetic data, we show that this downsampling method captures well the RR of actual NGS-read-reduced data (Fig. S5).

**FIG. 3.**
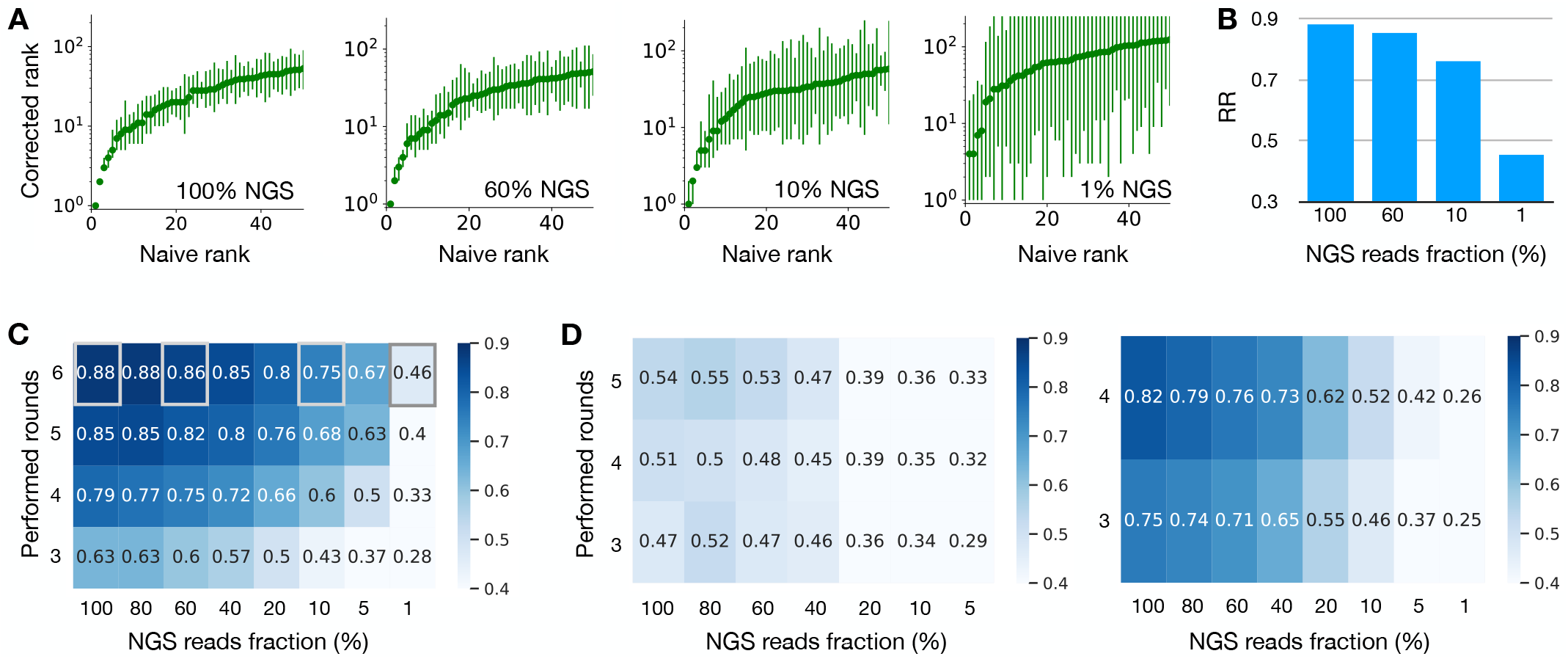
How the rank robustness depends on the experimental protocol. (A) Rank graphs for different NGS depths in Data-A (Table I). Different NGS-depth data are generated using downsampling (Materials and Methods). *x*% means the dataset where the number of NGS reads per round is reduced to *x*% (100% is the original dataset). (B) RR for the rank graphs in the panel A. Note that RR is higher than 0.7 even with the 10% NGS-depth. (C) The heat map showing RR for various NGS depths and performed rounds in Data-A. RR is larger than 0.7 for the data with (i) the 4 performed rounds with the NGS depth larger than or equal to 40% or with (ii) the 5 performed rounds with the NGS depth larger than or equal to 20%. This indicates that the data quality was already high with less experimental efforts. The four grey squares correspond to the four rank graphs in the panel A, respectively. (D) The same graphs as the panel C, but for different datasets. Data-E is used in the left panel, where RR is low and more NGS and/or screening rounds would be useful. Data-B is used in the right panel, where RR takes high values and seems to saturate in NGS depths. Further experimental efforts would probably not be necessary in this dataset.

### Pre-processing of Data-C and Data-D

In their original datasets, Data-C and -D contain a large number of variants whose total counts are very low (but not zero). In order to speed up the analysis and make the analysis more robust we removed the variants whose total counts are smaller than 1000 (Data-C) and than 100 (Data-D). The NGS depth and the number of unique variants shown in Table I are after this preprocessing.

## DATA AVAILABILTIY

All data analyzed in this article (Table I) are publicly available except for the random-peptide inserted library used for Fig. S1. This library will be deposited in a public database upon publication of this article.

## CODE AVAILABILITY

A Python implementation of ACIDES will be available on GitHub upon publication of this article.

## ACKNOWLEDGMENTS

The authors would like to thank A. Rubin and D. Fowler for kindly providing them with datasets (Data-A and Data-B) and a working code for enrich2. The authors also would like to thank L. C. Byrne and T. Mora for useful comments and discussions, M. Desrosiers and C. Robert for their technical assistance with the production of plasmids and viral vectors, and O. Marre for facilitating and initiating the collaboration and helpful discussions. This work was supported by ERC Starting Grant (REGENETHER 639888), European Research Council (ERC) Horizon 2020 Framework Programme Project: 863214 – NEUROPA, UNADEV, the Institut National de la Santé et de la Recherche Médicale (INSERM), Sorbonne Université, The Foundation Fighting Blindness, Agence National de Recherche (ANR) RHU Light4Deaf, LabEx LIFESENSES (ANR-10-LABX-65), IHU FORe-SIGHT (ANR-18-IAHU-01), and JSPS KAKENHI Grant Number 22K17994.

## Supplementary information for

**FIG. S1.**
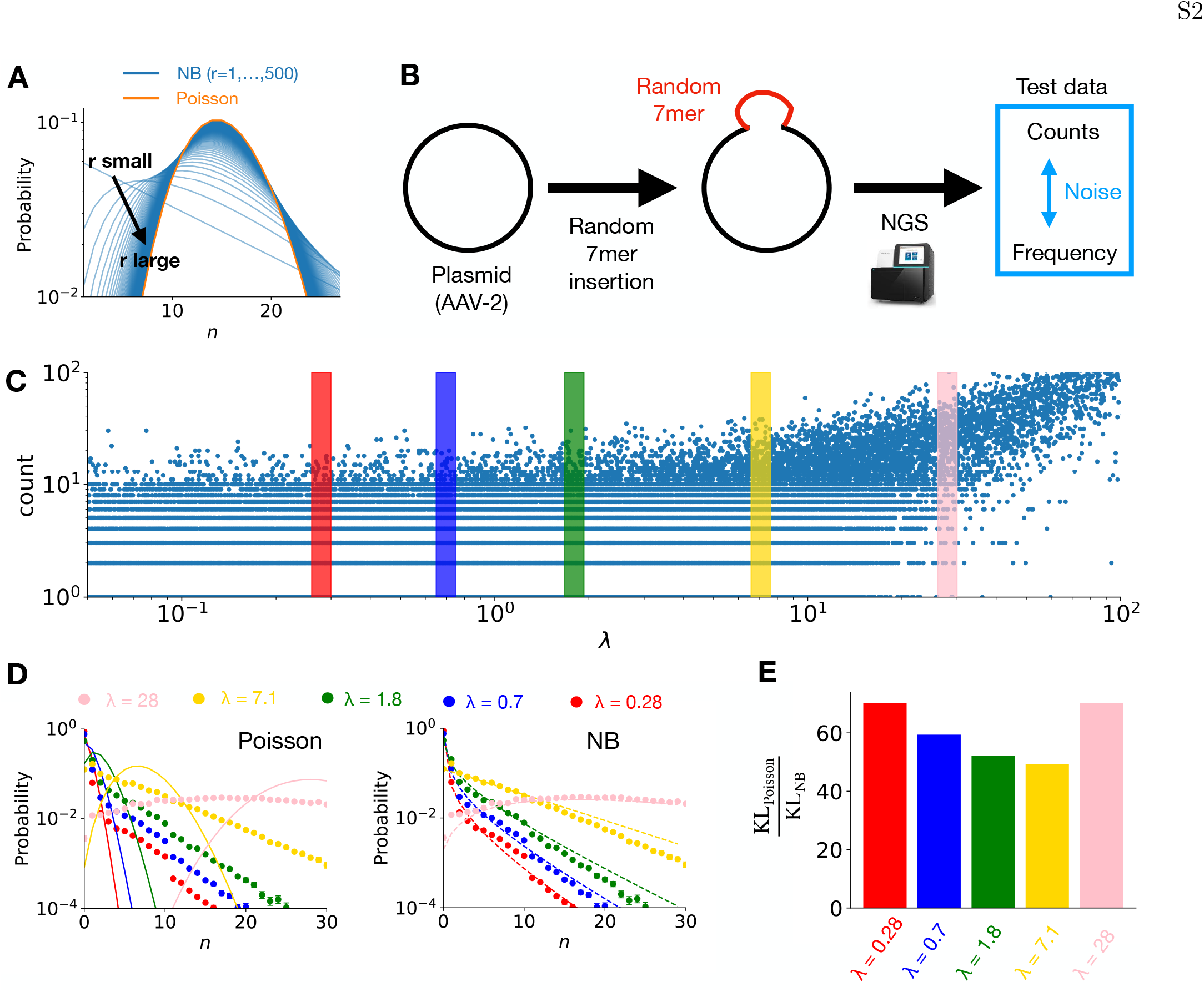
Negative binomial model accounts for NGS count noise better than Poisson model. (A) The poisson distribution (orange) and the negative binomial distribution (tableau blue) with the expected value *λ* = 15. The dispersion parameter *r* for the negative binomial distribution is set to 1, 2, …, 500. The negative binomial distribution generalizes the Poisson distribution, allowing for large variances by decreasing *r*. It converges to the Poisson distribution in the large *r* limit. (B) In order to test the predictive ability of the negative binomial distribution, we performed the following experiment. Using a random peptide (of size 21 corresponding to a 7mer) as a barcode, we first barcoded a plasmid extracted from adeno-associated virus 2 (AAV2) wild type. The obtained 7-mer inserted library was then sent to NGS facility and the corresponding barcoded region was sequenced. Since these 21 nucleotides of barcode are randomly and independently generated, we can use a position weight matrix model to predict the frequency of each variant in the sample. Comparing the predicted frequency with the actual NGS reads, we investigate the noise distribution of NGS counts. (C) The graph showing the obtained counts (*n*) against the predicted frequencies multiplied by the total NGS reads (*λ*), where each point corresponds to a variant. We observe that the counts are largely dispersed. (D) The probability distribution of counts for a fixed value of *λ* together with the model predictions by Poisson distribution (left) and by the negative binomial distribution (right). The probability distribution is estimated in the following way: (i) picking up all the variants within 5 different colored rectangles in the panel (C). (ii) Using the variants corresponding to each rectangle, we then make a histogram of counts, which is plotted in the panel (D) as dots. For the Poisson prediction, we simply use the Poisson distribution with the mean *λ* = 0.28, 0.7, 1.8, 7.1, 28. For the negative binomial prediction, for each value of *λ*, we infer the dispersion parameter *r* via a maximum likelihood inference and fitted a power-law function *r* = *βλ*^*α*^ to the obtained estimations. The result is *r* = 0.21*λ*^0.744^. Using this relation, the negative binomial distribution is then plotted for each *λ* = 0.28, 0.7, 1.8, 7.1, 28. (E) Comparison of predictive abilities between the poisson model and the negative binomial model. To quantify the predictive ability of each model, we use Kullback-Leibler divergence (KL) defined as KL = ∑_*n*_ *P*_data_(*n*) log(*P*_data_(*n*)*/P*_model_(*n*)). The ratio between KL for the poisson model and KL for the negative binomial model is plotted for each value of *λ*. KL_Poisson_ itself is 0.37, 0.59, 0.87, 1.59, 3.29 for *λ* = 0.28, 0.7, 1.8, 7.1, 28, while KL_NB_ is 0.0053, 0.0099, 0.017, 0.032, 0.047.

**FIG. S2.**
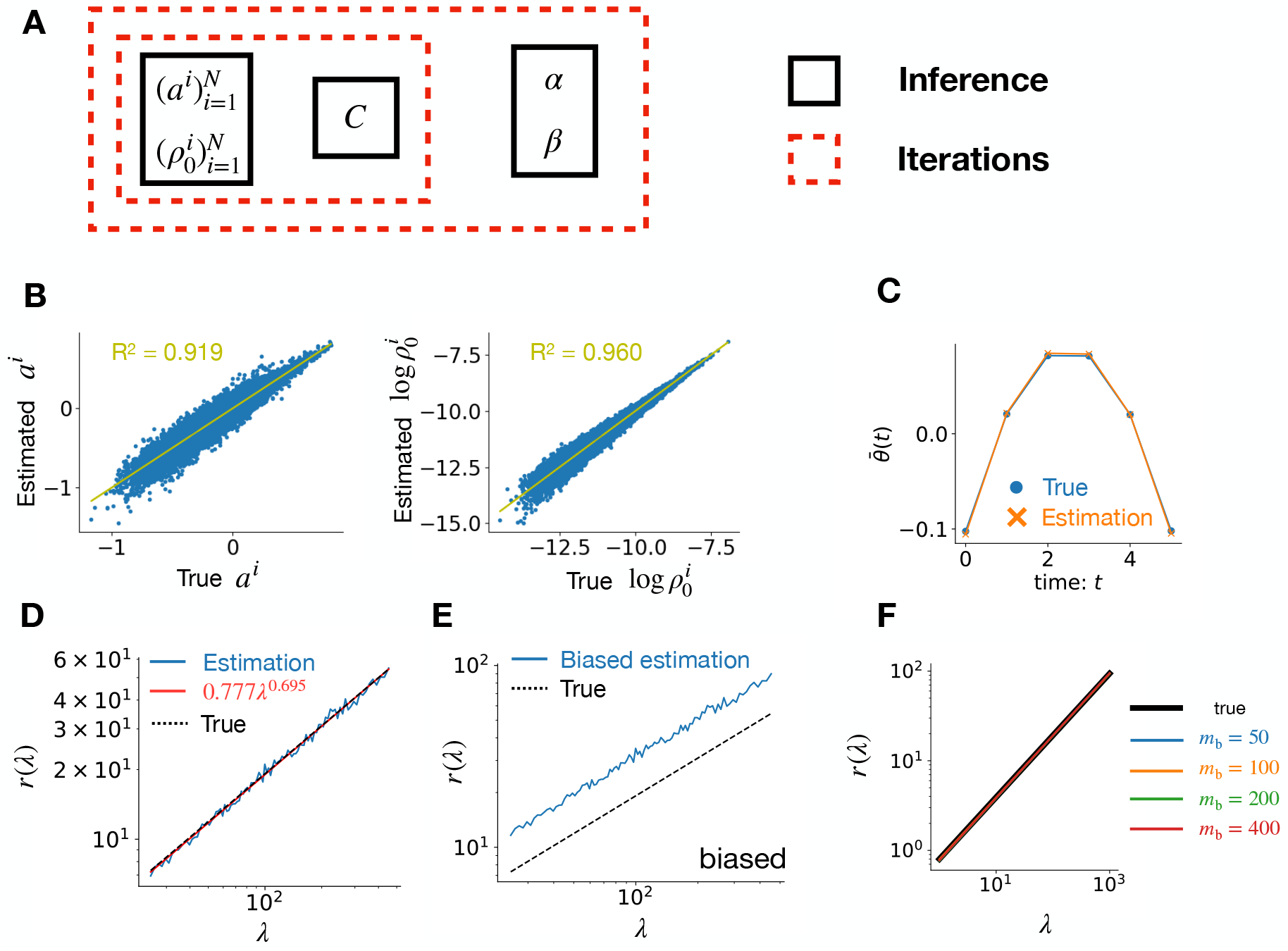
Inference algorithm and synthetic teacher-student examples. (A) graphical illustration of the inference algorithm with its double loop structure. The internal loop accounts for the parameters of the exponential model (see Materials and Methods) and iterates between the inference of 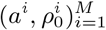 and *C*, while the external loop iterates between this internal loop and the inference of the negative binomial parameters (*α, β*). For the internal loop, we first infer 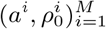 for a fixed *C, α, β* by maximizing the likelihood function (2). *C\** is then calculated from the obtained 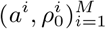 via 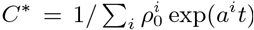 The linearly increasing part of log *C*^*\**^ is next subtracted as log *C*^*\**^ − (*x*^*\**^*t* + *y*^*\**^), where *x*^*\**^, *y*^*\**^ = argmax_*x,y*_ ∑_*t*_ [log *C* − *xt* − *y*]^2^ (fixing a gauge). The obtained quantity is log *C* for the next iteration. For the external loop, to obtain (*α, β*), we first infer the dispersion parameter *r* for a list of different values of *λ*. To do so, for a fixed value of *λ*, we select a set of index *i* and the time *t* by the condition 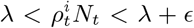 (with a small parameter *E*). Only using these *i* and *t*, we then maximize the likelihood function (2) and determine the value of *r*(*λ*). After obtaining the function *r*(*λ*) for several values of *λ*, we fit a linear function *r*(*λ*) = *αλ*^*β*^ to it and determine *α* and *β*. (B,C,D) Ground truth comparisons for the synthetic data-rich dataset (Fig. 1E) after 30th iterations demonstrates that the algorithm can recover the generating parameters. In the panel B, we plot the estimated parameters 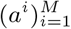 (left) and 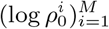 (right) against their ground truths. The coefficient of determination *R*^2^ is also shown. In the panel C, the normalization coefficient 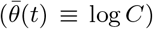 is plotted together with its ground truth. In the panel D, the estimated *r*(*λ*) with a fitted line *βλ*^*α*^ and its ground truth are shown. (E) While estimating *r*(*λ*) for a fixed *λ*, maximizing the likelihood function (2) results in a biased estimation as shown in the panel E. For fixing this, we generate a synthetic data probe using the current estimation of *α*_0_ and *β*_0_ with 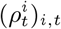 and use it to unbias the *r* estimation. More precisely, denoting the biased estimation by *r*_bias_(*λ*) (panel E) and the estimation of the *r* in the probe by *r*_1_(*λ*), the unbiased estimator plotted in the panel D is obtained as 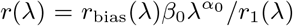. (F) To determine *α* and *β*, we use only representative variants, as described in Materials and Methods. For the representative variants, we select the variants whose total counts are larger than a threshold value *m*_b_. In (F), by using the synthetic data, we show that the inference results (*α* and *β*) are robust against the change of this parameter *m*_b_.

**FIG. S3.**
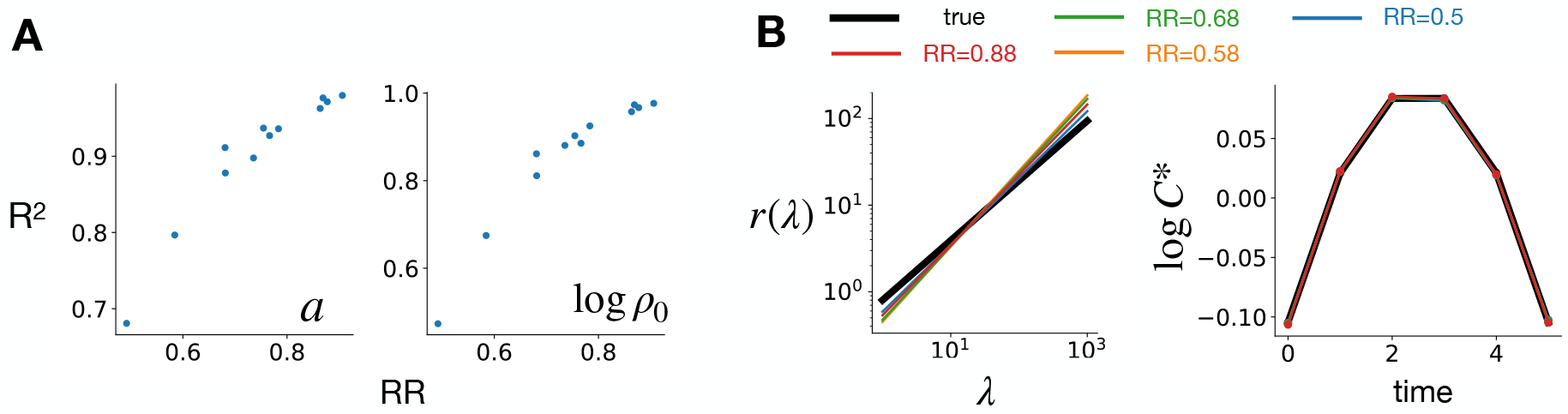
RR offers a proxy for the accuracy of the estimated scores. Here we use synthetic datasets, ranging from data-poor to data-rich regimes, to show that the empirical quantity RR correlates with our ability to recover the true values of scores *a*^*i*^ and initial frequencies 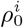. For this, we generate synthetic datasets with different values of total NGS reads *N* and number of variants *M* : (*N, M*) = (10^7^, 5 ×10^5^); (8 ×10^6^, 5× 10^5^); (6 ×10^6^, 5 ×10^5^); (4 ×10^6^, 5 ×10^5^); (2 ×10^6^, 5 ×10^5^); (10^6^, 5 ×10^5^); (10^7^, 10^6^); (8× 10^6^, 10^6^); (4 ×10^6^, 10^6^); (2× 10^6^, 10^6^); (10^6^, 10^6^). In these datasets, the parameters (*α, β*) are the same as those used in Fig.1E, F. (A) Coefficients of determination *R*^2^ between inferred and true values for *a*^*i*^ (left) and 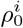 (right) are plotted against RR. This demonstrates the correlation between *R*^2^ and RR. (B) The inference of *r*(*λ*) and *C*^*\**^ is robust across all synthetic datasets.

**FIG. S4.**
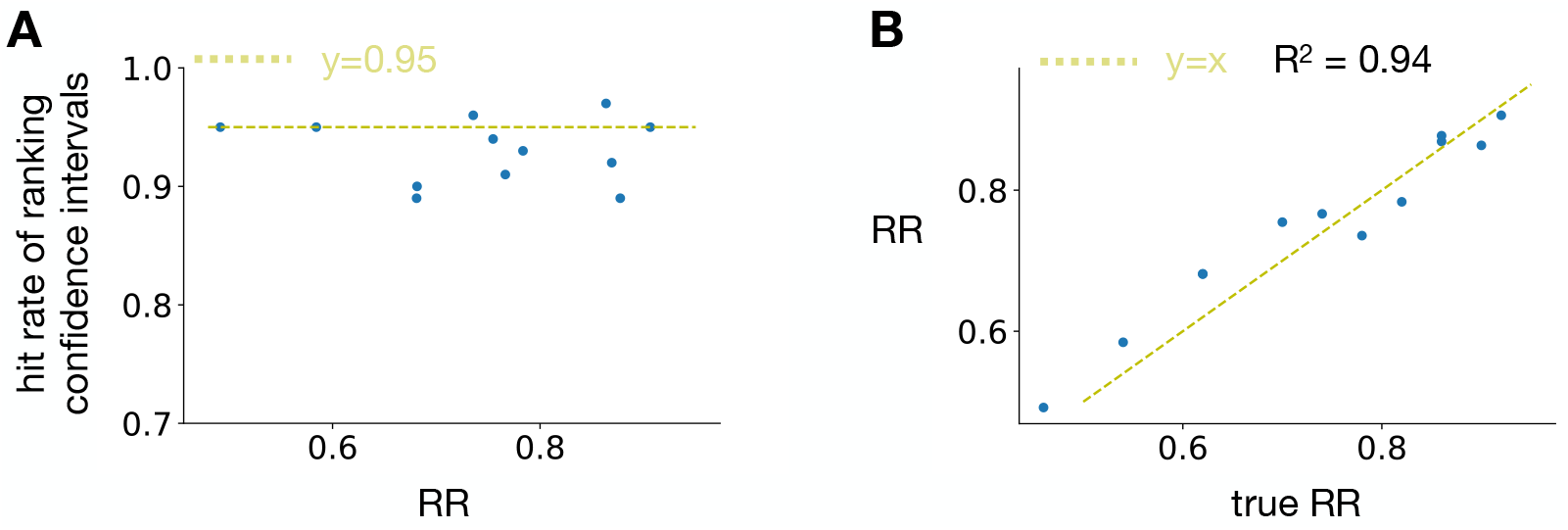
Ranking CI and rank robustness are accurately estimated also in the data-poor regime. A) For all the synthetic datasets introduced in Fig. S3. hit rates of the confidence interval of ranking graphs are plotted against their RR, where the hit rate is defined as the number of true ranking (red crosses in Fig.1E, F for example) that are within the 95%- confidence intervals (green lines in Fig.1E, F), divided by 50. The hit rates fluctuate around 0.95 as expected, demonstrating the quality of our estimation of ranking CI. B) Rank robustness (RR) estimated from inferred parameters is plotted against the true value (ground truth) for the synthetic datasets. The obtained high coefficient of determination shows that ACIDES can estimate RR also in the data-poor regime.

**FIG. S5.**
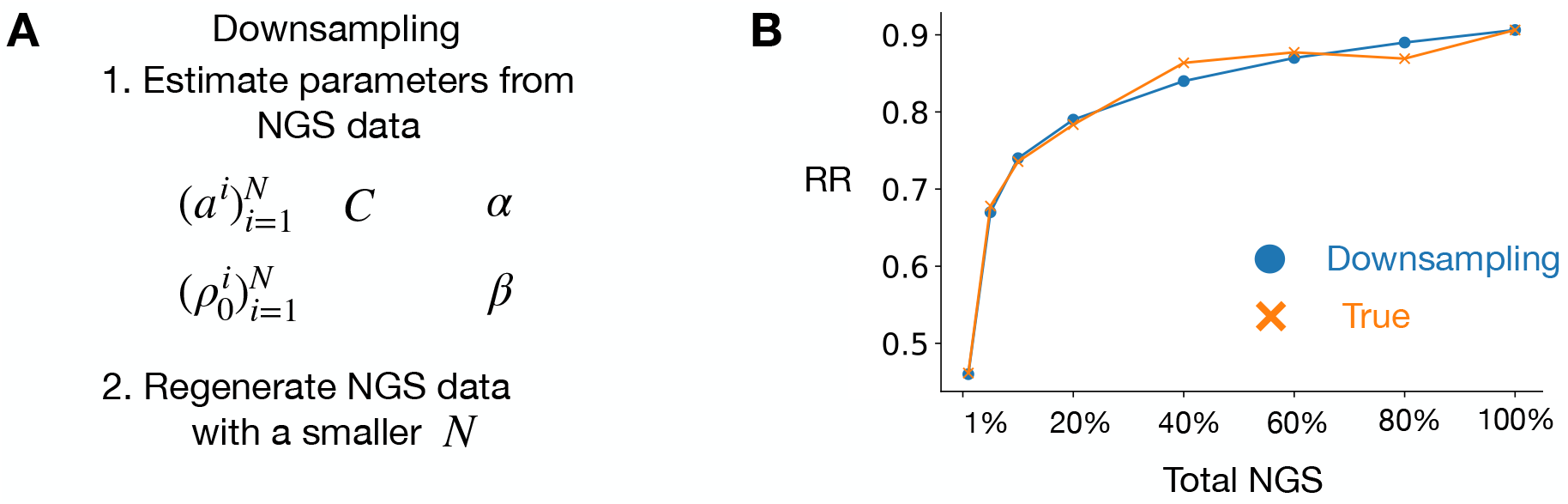
Downsampling NGS counts. (A) For a given dataset of a DE experiment, we downsample the data, *i*.*e*., create a synthetic dataset that corresponds to the dataset of the same DE experiments but with smaller values of the total NGS reads. For this, we first estimate the parameters of ACIDES from the dataset. We then generate synthetic NGS data using the likelihood function (2) with smaller numbers of the total NGS reads. For example, if we downsample the data to 40%, we set *N*_*t*_ to be 0.4*N*_*t*_. We then estimate RR using ACIDES for this downsampled dataset without reinferring *α, β*. (B) We show the validity of this down sampling method on the synthetic data. The parameters for the synthetic data are (*N*_*t*_, *M*) = (10^7^, 5×10^5^), (*α, β*) = (0.69, 0.8) and 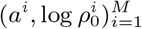 generated from the normal distribution with the expected values (-1,1) and the standard deviation (0.25, 1). We plot RRs obtained from this downsampling method (blue circles) and from a standard sampling method (orange crosses) as a function of the total number of NGS (where 100% means the original data). Here the standard sampling method means using ACIDES directly on the dataset with the total number of NGS 0.01*xN*_*t*_, where *x* is the percentage of the total NGS reads (*x*-axis in the panel B). We observe that our downsampling method estimates well the RR.

**FIG. S6.**
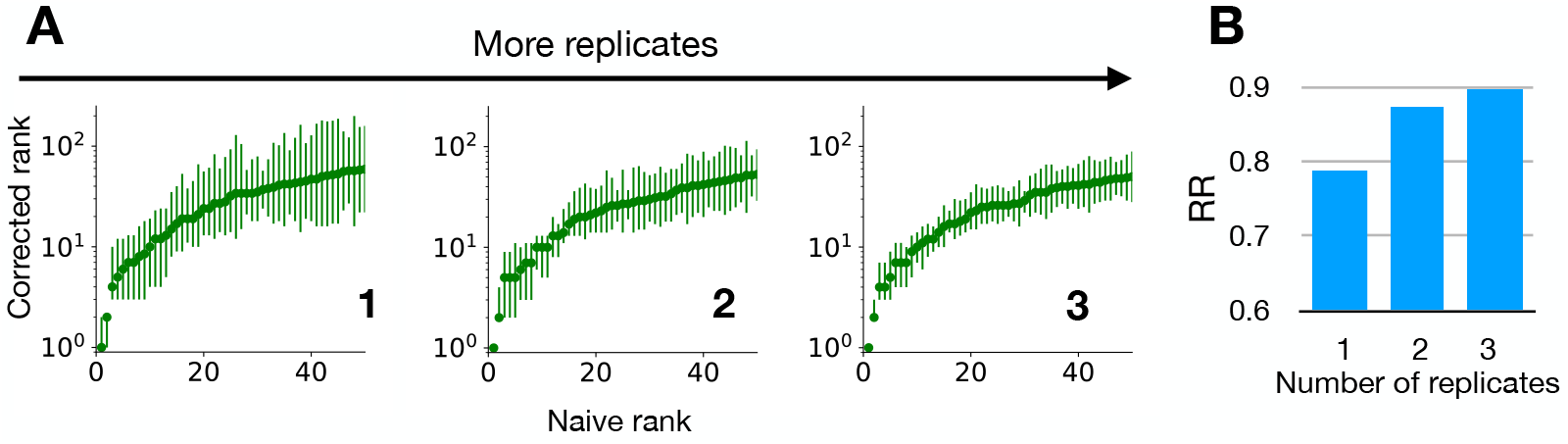
Multiple replicates can be combined to increase RR. We use the first 4 rounds of Data-A (so that RR is relatively small for a single replicate), and perform ACIDES for each replicate. (A) Ranking graphs for one (left), two (middle), three (right) replicates. To combine variant scores of two replicates (denoted by *a*_1_, *a*_2_ with standard deviation *δa*_1_, *δa*_2_), we use 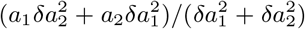 for the combined score and 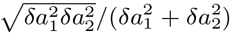 for the combined standard deviation. (B) RR for the three cases.

**FIG. S7.**
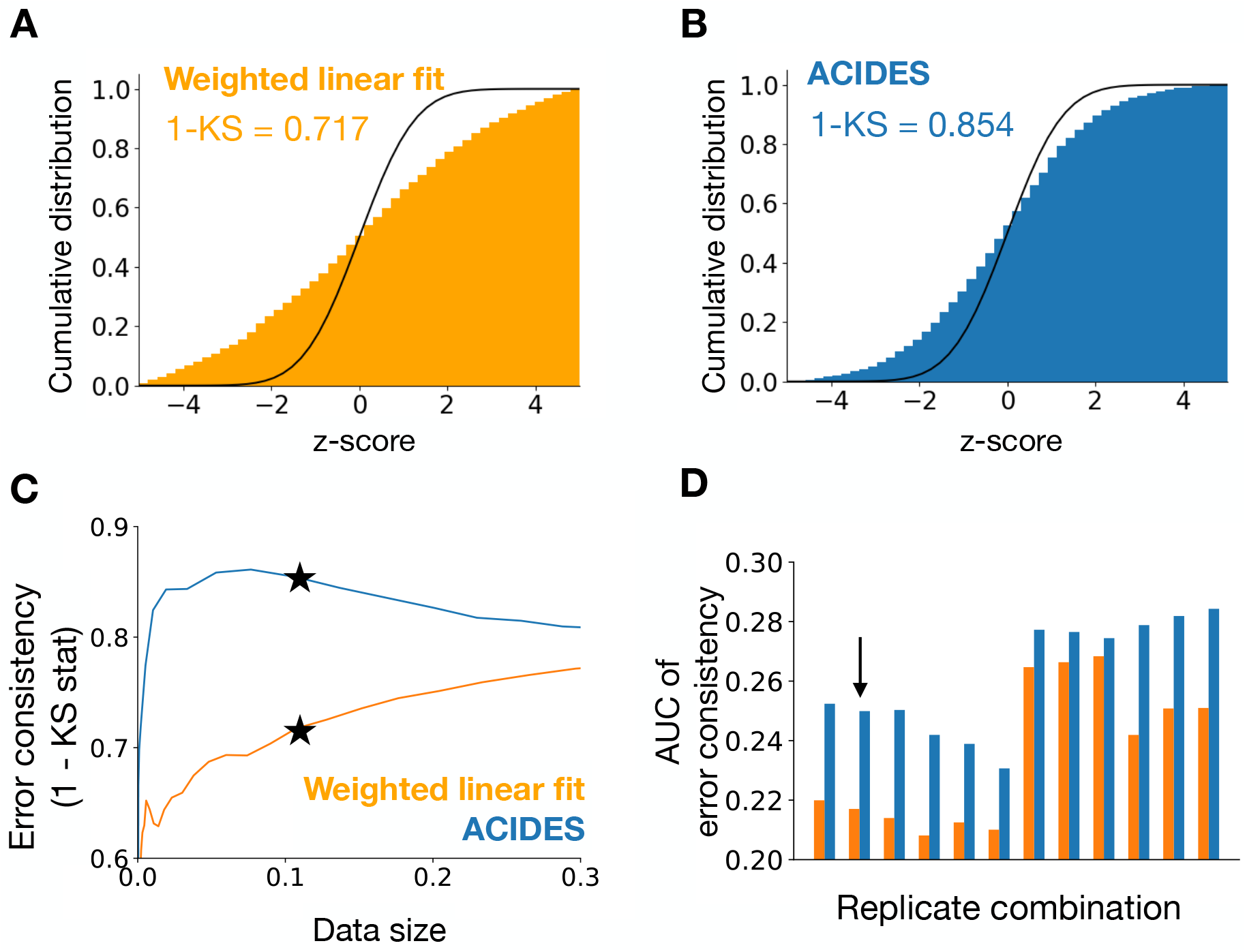
ACIDES outperforms previous methods in the estimation of score’s statistical errors. Using replicates in Data-A and Data-B (Table 1), we study the consistency of error bars in ACIDES and in *Enrich2*. Denoting by *a*_1_, *a*_2_ the scores of a variant in replicate 1 and 2 (similarly by *δa*_1_, *δa*_2_ the standard deviations of the scores), we study the following quantity 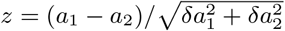 and compute the histogram of this quantity over different variants. Under the assumption that both scores are distributed following the normal distribution, this obtained histogram is approximated by the standard normal distribution. (A,B) Cumulative distributions of the histograms for *Enrich2* (A) and ACIDES (B) together with the cumulative standard normal distribution. We use 1 −Kolmogorov–Smirnov (KS) statistics (the maximum distance between two distributions measured in the y-direction) to quantify the distance between the histogram and the normal distribution. (C) To study 1 −KS more systematically, we introduce a threshold for the score statistical errors (Materials and Methods) by which we reduce the amount of data. For each fraction of the data, we estimate 1 −KS and plot them in the panel C. The stars in the panel correspond to panels A and B. (D) Finally, using all possible combinations of technical replicates in Data-A and Data-B, we compare ACIDES and *Enrich2*. We compute the area under curve (AUC) of 1− KS (the panel C) for all combinations. ACIDES always shows better performance than *Enrich2*. The arrow in the panel D indicates the replicate combination used in the panels A-C.

## Notes

### Competing Interest Statement

The authors have declared no competing interest.

### Summary of Updates

The title, the abstract and Figure 1 were revised.

## References

[1] Frances H Arnold. Design by directed evolution. Accounts of chemical research 31, 125 (1998).

[2] Philip A Romero and Frances H Arnold. Exploring protein fitness landscapes by directed evolution. Nature reviews Molecular cell biology 10, 866 (2009).

[3] Michael S Packer and David R Liu. Methods for the directed evolution of proteins. Nature Reviews Genetics 16, 379 (2015).

[4] Keqin Chen and Frances H Arnold. Tuning the activity of an enzyme for unusual environments: sequential random mutagenesis of subtilisin E for catalysis in dimethylformamide. Proceedings of the National Academy of Sciences 90, 5618 (1993).

[5] Nicholas J Turner. Directed evolution drives the next generation of biocatalysts. Nature chemical biology 5, 567 (2009).

[6] Olga Khersonsky Tawfik and Dan S. Enzyme Promiscuity: A Mechanistic and Evolutionary Perspective. Annual Review of Biochemistry 79, 471 (2010). PMID: 20235827.

[7] Robert E. Hawkins, Stephen J. Russell, and Greg Winter. Selection of phage antibodies by binding affinity: Mimicking affinity maturation. Journal of Molecular Biology 226, 889 (1992).

[8] Eric T Boder, Katarina S Midelfort, and K Dane Wittrup. Directed evolution of antibody fragments with monovalent femtomolar antigen-binding affinity. Proceedings of the National Academy of Sciences 97, 10701 (2000).

[9] Luca Perabo, Hildegard Büning, David M Kofler, Martin U Ried, Anne Girod, Clemens M Wendtner, Jörg Enssle, and Michael Hallek. In vitro selection of viral vectors with modified tropism: the adeno-associated virus display. Molecular Therapy 8, 151 (2003).

[10] Narendra Maheshri, James T Koerber, Brian K Kaspar, and David V Schaffer. Directed evolution of adenoassociated virus yields enhanced gene delivery vectors. Nature biotechnology 24, 198 (2006).

[11] Stefan Michelfelder and Martin Trepel. Adeno-associated viral vectors and their redirection to cell-type specific receptors. Advances in genetics 67, 29 (2009).

[12] Deniz Dalkara, Leah C. Byrne, Ryan R. Klimczak, Meike Visel, Lu Yin, William H. Merigan, John G. Flannery, and David V. Schaffer. In Vivo–Directed Evolution of a New Adeno-Associated Virus for Therapeutic Outer Retinal Gene Delivery from the Vitreous. Science Translational Medicine 5, 189ra76 (2013).

[13] Jakob Körbelin, Timo Sieber, Stefan Michelfelder, Lars Lunding, Elmar Spies, Agnes Hunger, Malik Alawi, Kleopatra Rapti, Daniela Indenbirken, Oliver J Müller, Renata Pasqualini, Wadih Arap, Jürgen A Kleinschmidt, and Martin Trepel. Pulmonary Targeting of Adenoassociated Viral Vectors by Next-generation Sequencingguided Screening of Random Capsid Displayed Peptide Libraries. Molecular Therapy 24, 1050 (2016).

[14] Leah C Byrne, Timothy P Day, Meike Visel, Jennifer A Strazzeri, Cécile Fortuny, Deniz Dalkara, William H Merigan, David V Schaffer, and John G Flannery. In vivo–directed evolution of adeno-associated virus in the primate retina. JCI insight 5 (2020).

[15] Mohammadsharif Tabebordbar, Kim A. Lagerborg, Alexandra Stanton, Emily M. King, Simon Ye, Liana Tellez, Allison Krunnfusz, Sahar Tavakoli, Jeffrey J. Widrick, Kathleen A. Messemer, Emily C. Troiano, Behzad Moghadaszadeh, Bryan L. Peacker, Krystynne A. Leacock, Naftali Horwitz, Alan H. Beggs, Amy J. Wagers, and Pardis C. Sabeti. Directed evolution of a family of AAV capsid variants enabling potent muscle-directed gene delivery across species. Cell 184, 4919 (2021).

[16] https://www.nobelprize.org/prizes/chemistry/2018/summary/.

[17] Sam Behjati and Patrick S Tarpey. What is next generation sequencing? Archives of Disease in Childhood -Education and Practice 98, 236 (2013).

[18] Shawn E. Levy and Richard M. Myers. Advancements in Next-Generation Sequencing. Annual Review of Genomics and Human Genetics 17, 95 (2016). PMID: 27362342.

[19] Douglas M Fowler, Carlos L Araya, Sarel J Fleishman, Elizabeth H Kellogg, Jason J Stephany, David Baker, and Stanley Fields. High-resolution mapping of protein sequence-function relationships. Nature methods 7, 741 (2010).

[20] Ryan T. Hietpas, Jeffrey D. Jensen, and Daniel N. A. Bolon. Experimental illumination of a fitness landscape. Proceedings of the National Academy of Sciences 108, 7896 (2011).

[21] Douglas M Fowler and Stanley Fields. Deep mutational scanning: a new style of protein science. Nature methods 11, 801 (2014).

[22] Alexandre Melnikov, Peter Rogov, Li Wang, Andreas Gnirke, and Tarjei S Mikkelsen. Comprehensive mutational scanning of a kinase in vivo reveals substratedependent fitness landscapes. Nucleic acids research 42, e112 (2014).

[23] Lea M Starita, David L Young, Muhtadi Islam, Jacob O Kitzman, Justin Gullingsrud, Ronald J Hause, Douglas M Fowler, Jeffrey D Parvin, Jay Shendure, and Stanley Fields. Massively Parallel Functional Analysis of BRCA1 RING Domain Variants. Genetics 200, 413 (2015).

[24] Sebastian Matuszewski, Marcel E Hildebrandt, Ana-Hermina Ghenu, Jeffrey D Jensen, and Claudia Bank. A Statistical Guide to the Design of Deep Mutational Scanning Experiments. Genetics 204, 77 (2016).

[25] Nathan J Rollins, Kelly P Brock, Frank J Poelwijk, Michael A Stiffler, Nicholas P Gauthier, Chris Sander, and Debora S Marks. Inferring protein 3D structure from deep mutation scans. Nature genetics 51, 1170 (2019).

[26] Jörn M Schmiedel and Ben Lehner. Determining protein structures using deep mutagenesis. Nature genetics 51, 1177 (2019).

[27] Rupali P Patwardhan, Choli Lee, Oren Litvin, David L Young, Dana Pe’er, and Jay Shendure. High-resolution analysis of DNA regulatory elements by synthetic saturation mutagenesis. Nature biotechnology 27, 1173 (2009).

[28] Matthew S Rich, Celia Payen, Alan F Rubin, Giang T Ong, Monica R Sanchez, Nozomu Yachie, Maitreya J Dunham, and Stanley Fields. Comprehensive Analysis of the SUL1 Promoter of Saccharomyces cerevisiae. Genetics 203, 191 (2016).

[29] Olga Puchta, Botond Cseke, Hubert Czaja, David Tollervey, Guido Sanguinetti, and Grzegorz Kudla. Network of epistatic interactions within a yeast snoRNA. Science 352, 840 (2016).

[30] Carlos L. Araya, Douglas M. Fowler, Wentao Chen, Ike Muniez, Jeffery W. Kelly, and Stanley Fields. A fundamental protein property, thermodynamic stability, revealed solely from large-scale measurements of protein function. Proceedings of the National Academy of Sciences 109, 16858 (2012).

[31] Alan F Rubin, Hannah Gelman, Nathan Lucas, Sandra M Bajjalieh, Anthony T Papenfuss, Terence P Speed, and Douglas M Fowler. A statistical framework for analyzing deep mutational scanning data. Genome biology 18, 1 (2017).

[32] Justus M. Kebschull and Anthony M. Zador. Sources of PCR-induced distortions in high-throughput sequencing data sets. Nucleic Acids Research 43, e143 (2015).

[33] Katharine Best, Theres Oakes, James M Heather, John Shawe-Taylor, and Benny Chain. Computational analysis of stochastic heterogeneity in PCR amplification efficiency revealed by single molecule barcoding. Scientific reports 5, 1 (2015).

[34] Vladimir Potapov and Jennifer L Ong. sources of error in PCR by single-molecule sequencing. PloS one 12, e0169774 (2017).

[35] Simon Festing and Robin Wilkinson. The ethics of animal research. EMBO reports 8, 526 (2007).

[36] Leah Byrne, Timothy Day, Meike Visel, Deniz Dalkara, Valerie Dufour, Felipe Pompeo Marinho, William Merigan, Gustavo Aguirre, William Beltran, David Schaffer, and John Flannery. Directed Evolution of AAV for Efficient Gene Delivery to Canine and Primate Retina - Raw counts of variants from deep sequencing. Dryad, Dataset (2018). https://doi.org/10.6078/D1895R.

[37] Sébastien Boyer, Dipanwita Biswas, Ananda Kumar Soshee, Natale Scaramozzino, Clément Nizak, and Olivier Rivoire. Hierarchy and extremes in selections from pools of randomized proteins. Proceedings of the National Academy of Sciences 113, 3482 (2016).

[38] Douglas M. Fowler, Carlos L. Araya, Wayne Gerard, and Stanley Fields. Enrich: software for analysis of protein function by enrichment and depletion of variants. Bioin-formatics 27, 3430 (2011).

[39] Simon Anders and Wolfgang Huber. Differential expres-sion analysis for sequence count data. Nature Precedings pages 1 (2010).

[40] Davis J. McCarthy, Yunshun Chen, and Gordon K. Smyth. Differential expression analysis of multifactor RNA-Seq experiments with respect to biological variation. Nucleic Acids Research 40, 4288 (2012).

[41] Michael I Love, Wolfgang Huber, and Simon Anders. Moderated estimation of fold change and dispersion for RNA-seq data with DESeq2. Genome biology 15, 1 (2014).

[42] Maximilian Puelma Touzel, Aleksandra M Walczak, and Thierry Mora. Inferring the immune response from repertoire sequencing. PLOS Computational Biology 16, e1007873 (2020).

[43] Bilge E Ö ztürk, Molly E Johnson, Michael Kleyman, Serhan Turunç, Jing He, Sara Jabalameli, Zhouhuan Xi, Meike Visel, Valérie L Dufour, Simone Iwabe, Luis Felipe L Pompeo Marinho, Gustavo D Aguirre, José-Alain Sahel, David V Schaffer, Andreas R Pfenning, John G Flannery, William A Beltran, William R Stauffer, and Leah C Byrne. scAAVengr, a transcriptome-based pipeline for quantitative ranking of engineered AAVs with single-cell resolution. eLife 10, e64175 (2021).

[44] Jorge Fernandez-de Cossio-Diaz, Guido Uguzzoni, and Andrea Pagnani. Unsupervised Inference of Protein Fitness Landscape from Deep Mutational Scan. Molecular Biology and Evolution 38, 318 (2020).

[45] Zachary Wu, S. B. Jennifer Kan, Russell D. Lewis, Bruce J. Wittmann, and Frances H. Arnold. Machine learning-assisted directed protein evolution with combinatorial libraries. Proceedings of the National Academy of Sciences 116, 8852 (2019).

[46] Richard J Fox, S Christopher Davis, Emily C Mundorff, Lisa M Newman, Vesna Gavrilovic, Steven K Ma, Loleta M Chung, Charlene Ching, Sarena Tam, Sheela Muley, et al. Improving catalytic function by ProSAR-driven enzyme evolution. Nature biotechnology 25, 338 (2007).

[47] Philip A. Romero, Andreas Krause, and Frances H. Arnold. Navigating the protein fitness landscape with Gaussian processes. Proceedings of the National Academy of Sciences 110, E193 (2013).

[48] Jakub Otwinowski, David M. McCandlish, and Joshua B. Plotkin. Inferring the shape of global epistasis. Proceedings of the National Academy of Sciences 115, E7550 (2018).

[49] Frédéric Cadet, Nicolas Fontaine, Guangyue Li, Joaquin Sanchis, Matthieu Ng Fuk Chong, Rudy Pandjaitan, Iyanar Vetrivel, Bernard Offmann, and Manfred T Reetz. A machine learning approach for reliable prediction of amino acid interactions and its application in the directed evolution of enantioselective enzymes. Scientific reports 8, 1 (2018).

[50] Claire N Bedbrook, Kevin K Yang, J Elliott Robinson, Elisha D Mackey, Viviana Gradinaru, and Frances H Arnold. Machine learning-guided channelrhodopsin engineering enables minimally invasive optogenetics. Nature methods 16, 1176 (2019).

[51] Kevin K Yang, Zachary Wu, and Frances H Arnold. Machine-learning-guided directed evolution for protein engineering. Nature methods 16, 687 (2019).

[52] Yuting Xu, Deeptak Verma, Robert P Sheridan, Andy Liaw, Junshui Ma, Nicholas M Marshall, John McIntosh, Edward C Sherer, Vladimir Svetnik, and Jennifer M Johnston. Deep dive into machine learning models for protein engineering. Journal of chemical information and modeling 60, 2773 (2020).

[53] Drew H Bryant, Ali Bashir, Sam Sinai, Nina K Jain, Pierce J Ogden, Patrick F Riley, George M Church, Lucy J Colwell, and Eric D Kelsic. Deep diversification of an AAV capsid protein by machine learning. Nature Biotechnology 39, 691 (2021).

[54] Claudia Bank, Ryan T Hietpas, Alex Wong, Daniel N Bolon, and Jeffrey D Jensen. A Bayesian MCMC approach to assess the complete distribution of fitness effects of new mutations: uncovering the potential for adaptive walks in challenging environments. Genetics 196, 841 (2014).

[55] Jakub Otwinowski. Biophysical Inference of Epistasis and the Effects of Mutations on Protein Stability and Function. Molecular Biology and Evolution 35, 2345 (2018).

[56] Luca Sesta, Guido Uguzzoni, Jorge Fernandez-de Cossio-Diaz, and Andrea Pagnani. AMaLa: Analysis of Directed Evolution Experiments via Annealed Mutational Approximated Landscape. International Journal of Molecular Sciences 22 (2021).

[57] Andrea Di Gioacchino, Jonah Procyk, Marco Molari, John S. Schreck, Yu Zhou, Yan Liu, Rémi Monasson, Si-mona Cocco, and Petr Šulc. Generative and interpretable machine learning for aptamer design and analysis of in vitro sequence selection. PLOS Computational Biology 18, 1 (2022).

[58] James T Koerber, Narendra Maheshri, Brian K Kaspar, and David V Schaffer. Construction of diverse adenoassociated viral libraries for directed evolution of enhanced gene delivery vehicles. Nature protocols 1, 701 (2006).

